# The long-term dynamics of the CA1 spatial code preserve the relative structure of contextual representation

**DOI:** 10.1101/2021.01.12.426395

**Authors:** Alexandra T. Keinath, Coralie-Anne Mosser, Mark P. Brandon

## Abstract

The hippocampus is thought to mediate episodic memory through the instantiation and reinstatement of context-specific cognitive maps. However, recent longitudinal experiments have challenged this view, reporting that most hippocampal cells change their tuning properties over days even in the same environment. Often referred to as *representational drift*, these dynamics raise multiple questions about the representational capacity and content of the hippocampal code. One such question is whether and how these dynamics impact the contextual code. To address this, here we imaged large CA1 populations over more than a month of daily experience as freely behaving mice participated in an extended geometric morph paradigm. We find that long-timescale changes in population activity occurred orthogonally to the representation of context in network space, allowing for consistent readout of contextual information across weeks. Together, these results demonstrate that long-timescale changes to the CA1 spatial code preserve the relative structure of contextual representation.

## Introduction

Hippocampal subregion CA1 represents a mixture of external and internal cues – including the shape of the environment^1^, visual landmarks^2,3^, objects^4^, task-relevant information^5^, and past experience^6–9^ – through changes to the spatial tuning properties of its principal cells^10^. Often collectively referred to as a *cognitive* or *hippocampal map*^11^, this code is hypothesized to support spatial and episodic memory by reinstating content at later times, for example by reinstantiating the same map across repeated visits to the same environment. However, recent longitudinal experiments in mice have challenged this view^12–16^. These studies report that the majority of cells change their spatial tuning properties, such as firing rates^13^ and field locations^14^, over a timescale of days, yielding maps of the same environment that, across time, are as different as maps between environments when assessed by various population-level measures. Despite these population-level dynamics, however, a minority of cells can maintain their spatial tuning properties across a months-long timescale under at least some circumstances.

These long-term dynamics raise a variety of questions about the representational capacity and content of the hippocampal code^17^, questions which might be resolved in multiple ways. On one hand, some suggest that these dynamics reflect the unavoidable biological realities of this code rather than its representational content^17–19^. That is, while the hippocampal code might serve a traditional mapping function, it is constrained by its biological instantiation. Because the content of this code must be inferred from inherently variable and plastic circuits, this account claims, its representation will *drift* over time, i.e. the representation will change even in the absence of changes to the content of the representation. On the other hand, it is possible that these dynamics instead faithfully reflect changes in the content of the representation. Although an environment remains the same across repeated visits in a spatial sense, each visit differs due to a variety of factors such as time, intervening experience, and idiosyncratic external cues which might elude the control of the experimenter. Given the responses of the CA1 spatial code to such diverse content on short timescales, it is thus plausible that the long-term dynamics are not a biological accident but instead accurately recapitulate the heterogenous dynamics of its content. Yet another possibility is that these dynamics point toward a necessary revision of function beyond a mapping framework, for example one that emphasizes the temporal structure over the spatial correlates of this code.

Central to the debate between these views is the relationship between the representational capacity and the long-term dynamics of the CA1 code. While prior work has informatively characterized long-term dynamics within the same environment and across two highly distinct environments^13–16^, such work is limited in its ability to speak to this relationship. Repeated recordings in the same environment characterize only effects occurring within that spatial context, whereas recording in two distinct environments has yielded representations which begin and remain orthogonal throughout, and therefore any changes to the relative representational structure cannot be characterized further.

To address this knowledge gap, we imaged large CA1 populations over more than a month of daily experience as freely behaving mice explored six differently-shaped environments in an extended geometric morph paradigm^8^. This paradigm elicited partially-correlated population-level maps, allowing us to characterize how the relative representational structure evolves over long timescales. We found that individual cells represent spatial context through heterogenous but stereotyped changes to their spatial tuning properties which led to a dynamic attractor-like population response. Characterizing the full representational structure of all sessions revealed that long-timescale changes in population activity occurred orthogonally to the representation of context in network space. Thus, despite continued representational changes over the course of the experiment, the relative representational structure of the environments was preserved, allowing for consistent readout of contextual information across weeks. Together these results demonstrate that long-timescale changes to the CA1 spatial code do not distort the landscape of contextual representation.

## Results

We recorded daily from large CA1 populations via calcium imaging (Fig. 1a) as mice freely explored open environments for 32 days in an extended version of a geometric morph paradigm (5 mice, 160 sessions total; Fig. 1b; Fig. S1). In this paradigm, mice were initially familiarized with two geometrically-distinct environments, and later tested in these environments as well as four deformed (morphed) versions of these environments spanning the shapespace between the two familiar environments. On each test day, activity was recorded in a single environment, after which the mice received additional unrecorded top-up experience in the remaining familiar environment(s). The test phase began with recording both familiar environments across two days (order randomized across mice). Next, activity was recorded in each morph environment (in a random order) over four days. Then the familiar environments were recorded again to bookend that morph sequence. This pattern was repeated for 32 testing days at which point 5 full bookended morph sequences were recorded. The order of unrecorded experience was randomized on each day. All sessions (and all unrecorded top-up experience) lasted 20 min per environment. Only one environment was recorded per day to reduce the risk of photobleaching.

**Figure 1.**
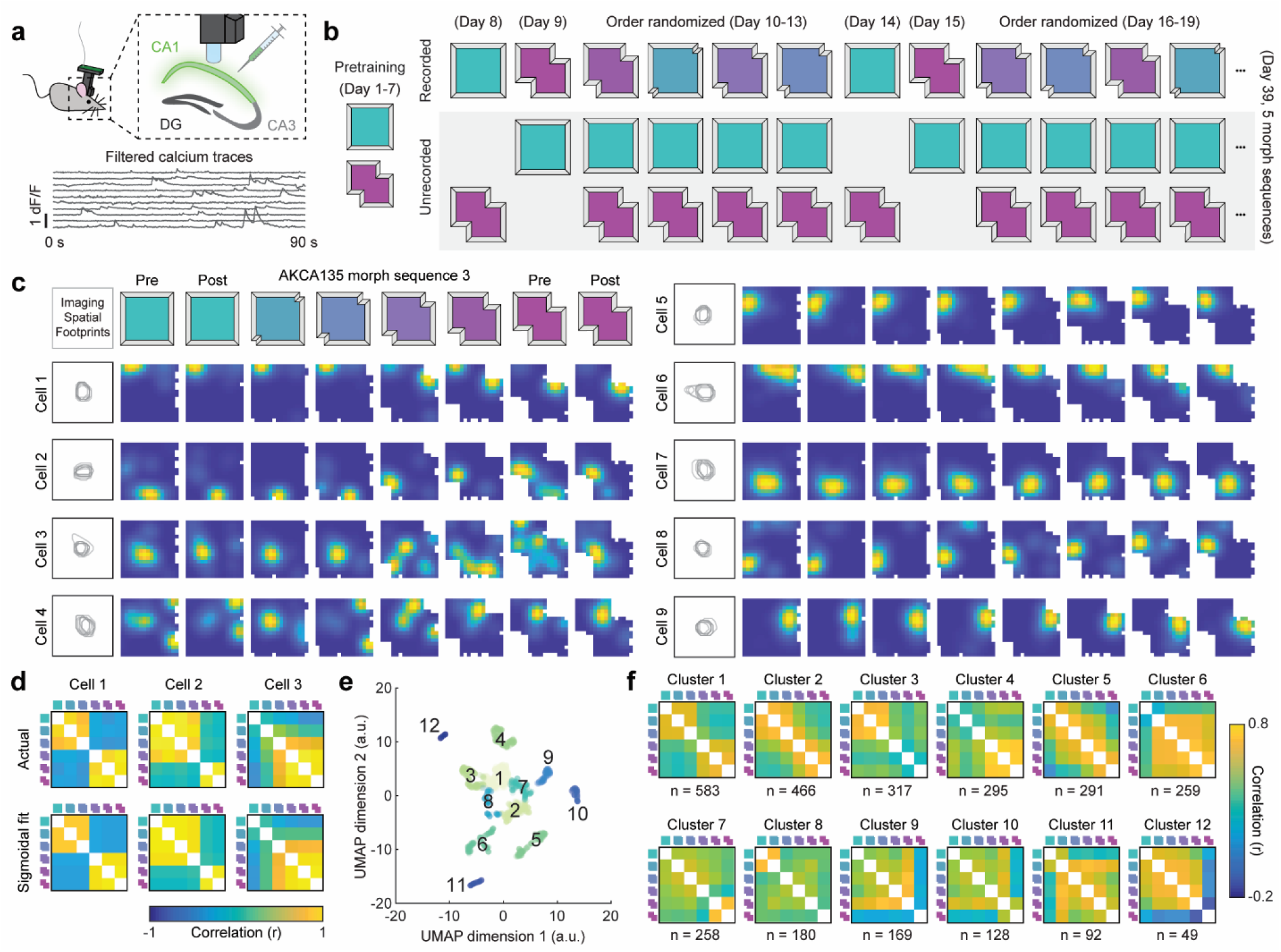
An adapted geometric morph paradigm can characterize the CA1 spatial code across extended experience. (**a**) Schematic of the miniscope recording procedure (top), as well as resulting calcium traces and firing rates inferred from second order autoregressive deconvolution (bottom). (**b**) Schematic of the behavioral paradigm. (**c**) Example of nine simultaneously recorded cells from mouse AKCA135 tracked across one morph sequence exhibiting a diversity of dynamics. Extracted spatial footprints for all 8 sessions shown left of rate maps. Maps normalized from zero (blue) to the peak (yellow) within each session. (**d**) Example actual contextual representational similarity matrices (RSMs) for a 6-day morph sequence for three cells (top), and the 5-parameter sigmoidal fit (bottom). (**e**) UMAP-embedding of all 3087 contextual RSMs based on their 5 sigmoidal fit parameters, clustered via k-medoids with k = 12. (**f**) Mean RSM for each cluster in (**e**) with number of RSMs in each cluster noted below.

We first characterized these data within each morph sequence. Following motion correction^20^, cells were segmented and calcium traces were extracted via constrained nonnegative matrix factorization^21,22^ (Fig. S1). The rising phase of transients was extracted from the filtered calcium traces (see *Methods*), and this binary vector was treated as the *firing rate* in all further analyses. Cell identity was tracked across recording sessions on the basis of mean imaging frame landmarks and the extracted spatial footprints (n = 4070)^23^.

We observed heterogenous but stereotyped responses at the level of individual cells (Fig. 1c). Some cells exhibited attractor-like properties in their spatial tuning, with either an abrupt transition in their preferred firing locations over the course of the morph sequence or the expression of a punctate field in only a contiguous subset of the shapespace. Other cells continued to fire at geometrically-similar locations across all environments spanning the shapespace. To explore this combination of heterogeneity and stereotypy, we extracted the contextual representational similarity matrix (RSM) across each 6-day morph sequence for all cells tracked across all 6 days. If a cell was tracked across multiple complete 6-day sequences, then it contributed multiple contextual RSMs, up to 5 RSMs per cell (3087 total RSMs). Next, we fit each of these with a 5-parameter sigmoidal model which could capture a large range of interpretable contextual dynamics such as gradual and abrupt transitions across contexts, asymmetric representations of subsets of the shapespace, and varying degrees of modulation (Fig. 1d; see *Methods*). Actual RSM fits resulted in significantly less error than fits to shuffled RSMs, and less error than a 5-parameter model in which RSM dynamics were instead described by two second-order polynomials (see *Methods*; Fig. S2), indicating that the sigmoidal model provided a better description of contextual RSM dynamics than would be expected by chance or from a more general functional form with equivalent degrees of freedom. We then reduced this 5-parameter sigmoidal description down to 2-dimensions via Uniform Manifold Approximation and Projection (UMAP), where RSMs appeared to group together (Fig. 1e). Unsupervised clustering of RSM similarity in this embedding via *k*-medoids into 12 clusters revealed that each cluster was defined by a particular pattern of contextual dynamics (Fig. 1f). The dominant clusters were defined by dynamics previously described in acute versions of this paradigm^8,9^: an abrupt transition in map similarity near the midpoint of the shapespace, high map similarity in one half of the shapespace but not the other, and a gradual change in map similarity across the shapespace. In addition to these dynamics, this procedure also identified smaller clusters of RSMs which were described by abrupt transitions in map similarity occurring at points other than the middle of the shapespace.

At a population level, these single-cell responses drove sigmoidal changes in mean rate map similarity across the shapespace, such that maps remained similar to the nearby familiar environment map, with an abrupt transition in similarity near the midpoint of the shapespace, similar to previous acute reports^8,9^ (Fig. 2a). In four of the five mice, the point of transition remained near the midpoint of the morph sequence across all five sequences; in one mouse it began near the midpoint of the first morph sequence, but gradually shifted over time (Fig. 2b), also similar to a previous report^8^. In all cases, the familiar and morph environments remained partially correlated with one another, though we observed a continues decorrelation across morph sequences such that the final morph sequence showed significantly more decorrelation than the first morph sequence in all mice (Fig. 2c). This gradual decorrelation might be a result of experience with the morph sequence itself or the ongoing daily unrecorded top-up experience in each of the familiar environments. Together, these results demonstrate that our adapted geometric morph paradigm replicates many phenomena observed within morph sequence in acute versions of this paradigm, and extends these findings to demonstrate that these within-sequence dynamics evolve with continued experience.

**Figure 2.**
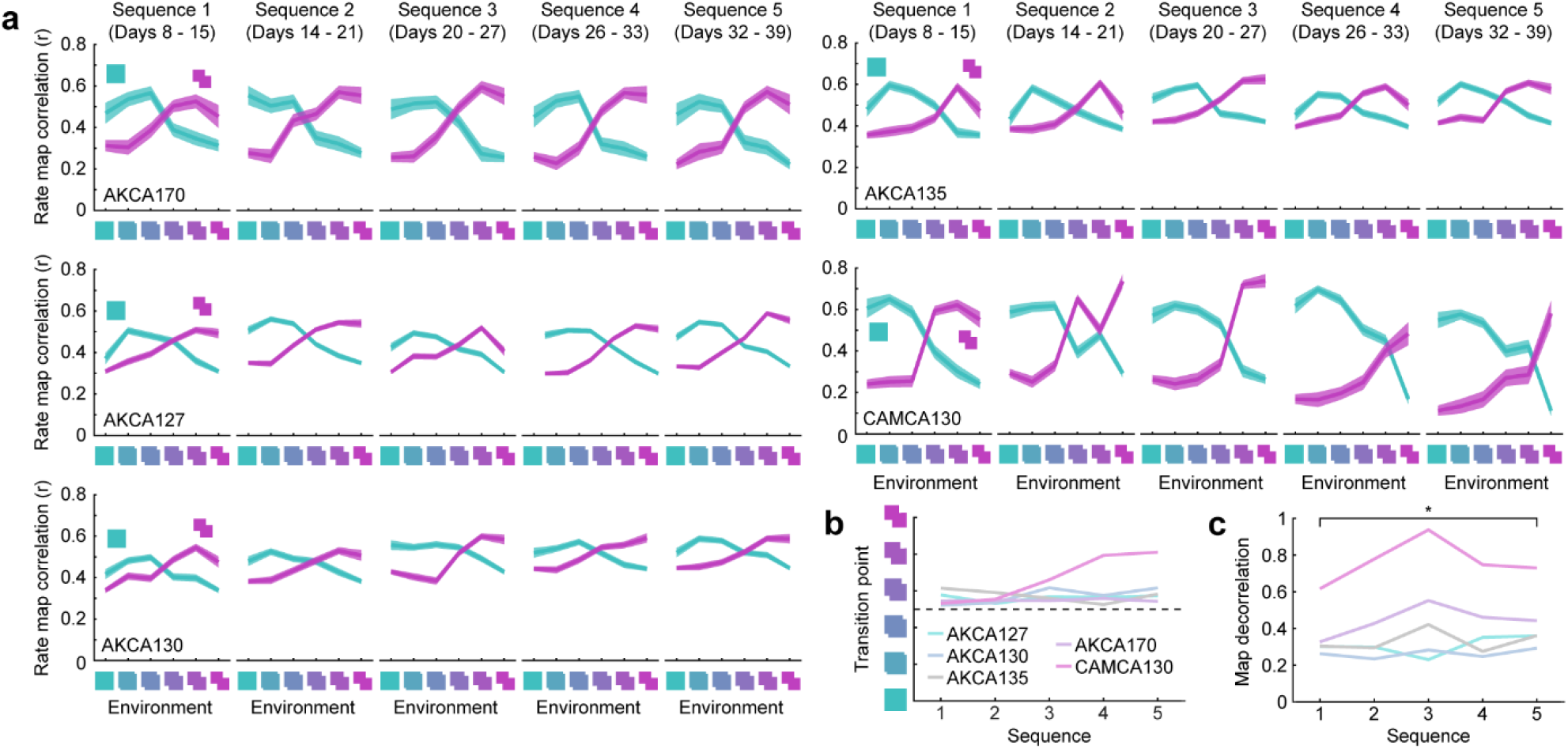
Within-sequence population morph dynamics evolve with extended experience. (a) Rate map correlations between each environment and the two familiar environments for all five eight-day bookended morph sequences for all five mice. Mouse name in lower left. Lines and shading denote mean ± 1 SEM across all cells whose within-session split-half reliability exceeding the 95^th^ percentile of a shuffled distribution for at least one of the compared sessions. (b) Transition point as a function of sequence number. Transition point was computed by first fitting the Fisher-transform of the two familiar environment rate map correlations with sigmoidal functions, and then determining their point of intersection. (**c**) Decorrelation between the familiar environment rate maps as a function of sequence number. Computed as the maximum difference between the two sigmoidal fits described in (**b**). Familiar environment rate maps were more decorrelated in the final sequence than the initial sequence (paired t-test: t(4) = 4.42, p = 0.0114). *p < 0.05

Next, we explored how population-level representational similarity changes across all 32 days of recording. To this end, we first computed the mean rate map similarity across all tracked cells for each pairwise comparison of sessions (Fig. 2a). These matrices exhibited structural features which generalized across mice: mean rate map correlations vary from high values for pairs of sessions close in time toward zero for sessions pairs separated further in time, with additional modulation by contextual similarity. To make this structure explicit, we then reduced this mean pairwise rate map similarity matrix to two dimensions via nonmetric multidimensional scaling (MDS), an unsupervised technique for visualizing high-dimensional structure in a digestible low-dimensional space that preserves the representational similarity between sessions as well as possible. The resultant embedding revealed two dimensions that strongly determined representational similarity in this paradigm: a contextual component and a drift component (Fig. 2b). These components were pronounced in every mouse. Quantification of the embedded representations revealed that the contextual and drift components defined nearly orthogonal dimensions in this 2D subspace, with absolute angular differences between context and drift dimensions neighboring 90° (see *Methods*; all angles: [83.9°, 88.0°, 91.1°, 95.4°, 98.8°]; Rayleigh’s test versus uniformity on the 0° to 180° range: p = 2.3e-3, z = 4.83). Estimations of the direction of drift from both familiar environments were similar to one another (absolute angular difference between drift direction estimates: [7.6°, 13.1°, 18.8°, 22.0°, 24.4°]). Similar results were observed when using population vector correlations as the measure of similarity between pairwise session comparisons (Fig. S3).

To situate these findings, we constructed a computational model in which individual CA1 cells were simulated as the rectified sum of spatially-tuned inputs modulated by drift, contextual group (grouping together each half of the shapespace), or the shape of the environment (Fig. 2c; see Methods). Next, we varied the dynamics of the drift input, such that drift accrued dependent on the specific environment, the context group, or simply as a function of time (global drift). Each drift dynamic resulted in a different stereotyped MDS embedding (Fig. 2d). Global drift yielded an embedding in which drift and contextual components were orthogonal to one another, and the direction of drift estimated from both familiar environments were consistent with one another (Fig. 2e). Contextual group-dependent drift also resulted in roughly orthogonal drift and contextual components; however, estimates of the direction of drift differed between the two familiar environments (Fig. 2e). Environment-specific drift yielded a qualitatively distinct inside-outward radial embedding in which each environment became more distinct over time, leading to roughly opposing estimates of the direction of drift between the familiar environments (Fig. 2e). Thus, only global drift reproduced essential characteristics of the representational structure of the CA1 spatial code in this paradigm.

If drift preserves the representational landscape of context in this paradigm, then we should be able to accurately predict contextual information across time, even on the basis of a restricted temporally-contiguous subset of the data. To address this possibility, we attempted to predict both context group identity (again grouping together each half of the shapespace) and the specific environment identity on the basis of population vector rate map similarity to each session in a contiguous 6-day predictor morph sequence (Fig. 3a). Only cells tracked across all six predictor sessions and the target session were included. In both cases, prediction accuracy remained above chance in all animals even 19+ days (>3 sequences) after the predictor sequence, resulting in significantly above-chance prediction accuracy at every epoch with ANOVAs indicating no significant differences between epochs (Fig. 3b,c). Together, these results provide additional evidence that drift preserves the relative representational structure of context, allowing accurate readout of contextual information across weeks-long timescales in this paradigm.

**Figure 3.**
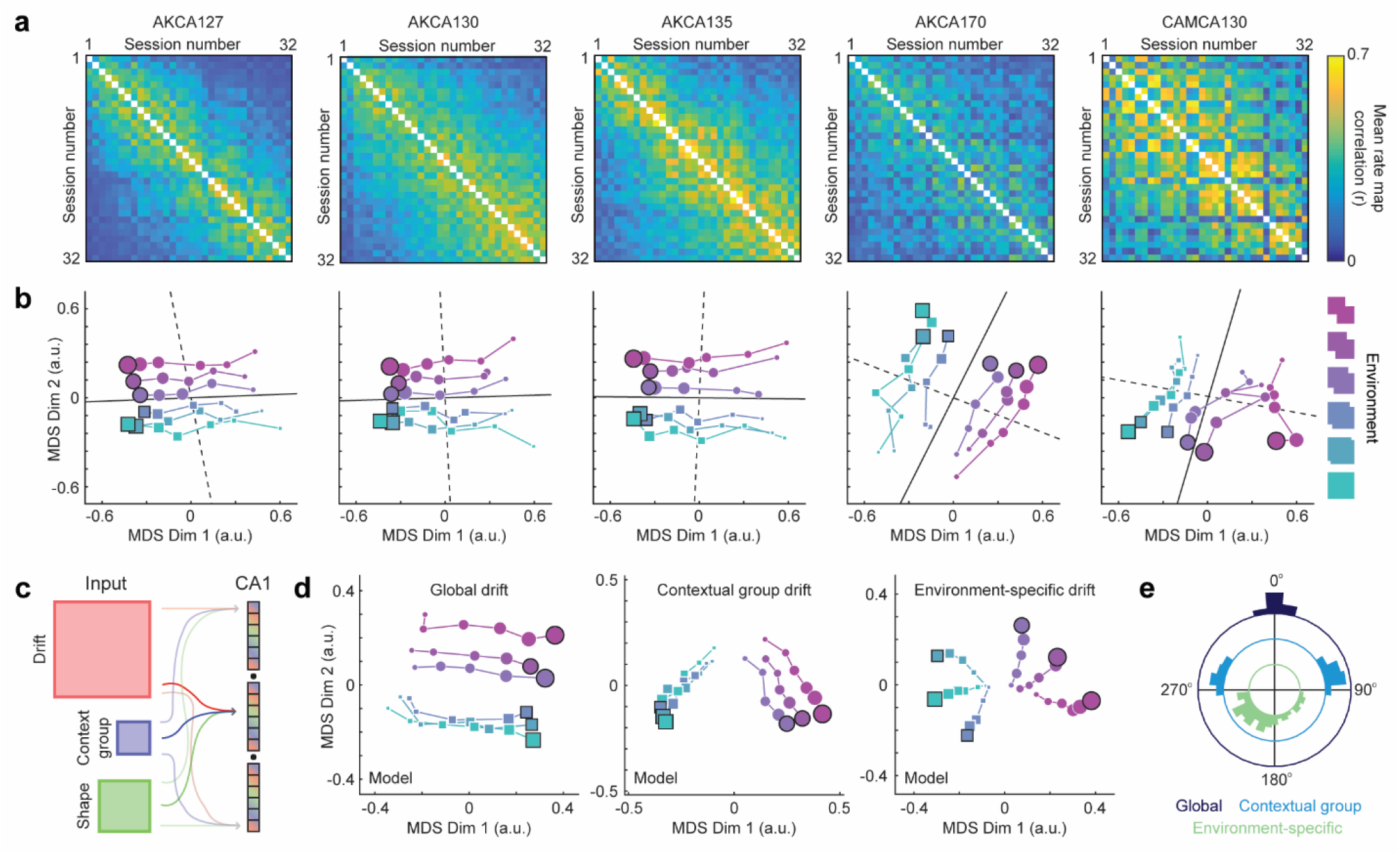
Dimensionality reduction reveals distinct temporal and contextual components governing the CA1 spatial code. (**a**) Population-level representational similarity matrices across all recording sessions for all five mice. Each RSM is computed by taking the mean rate map correlation across all tracked cells for each pairwise comparison of sessions. Mouse name at the top of each column. (**b**) Session similarity structure when embedded in a two-dimensional space via nonmetric multidimensional scaling (MDS). Dot size indicates session number, with earliest sessions indicated by the smallest dots and latest sessions indicated by the largest dots. Final sessions outlined in black. Note that estimated drift (solid black line) and context (dashed black line) dimensions are nearly orthogonal in this embedding (see Methods). (**c**) CA1 populations were modeled as a combination of spatial inputs which were modulated by drift, contextual group, and the shape of the environment. (**d**) Example MDS embeddings of modeled populations for different drift dynamics. (**e**) Distributions of the angular difference between the direction of drift estimated from both familiar environments for global, contextual group, and environment-specific drift dynamics.

**Figure 4.**
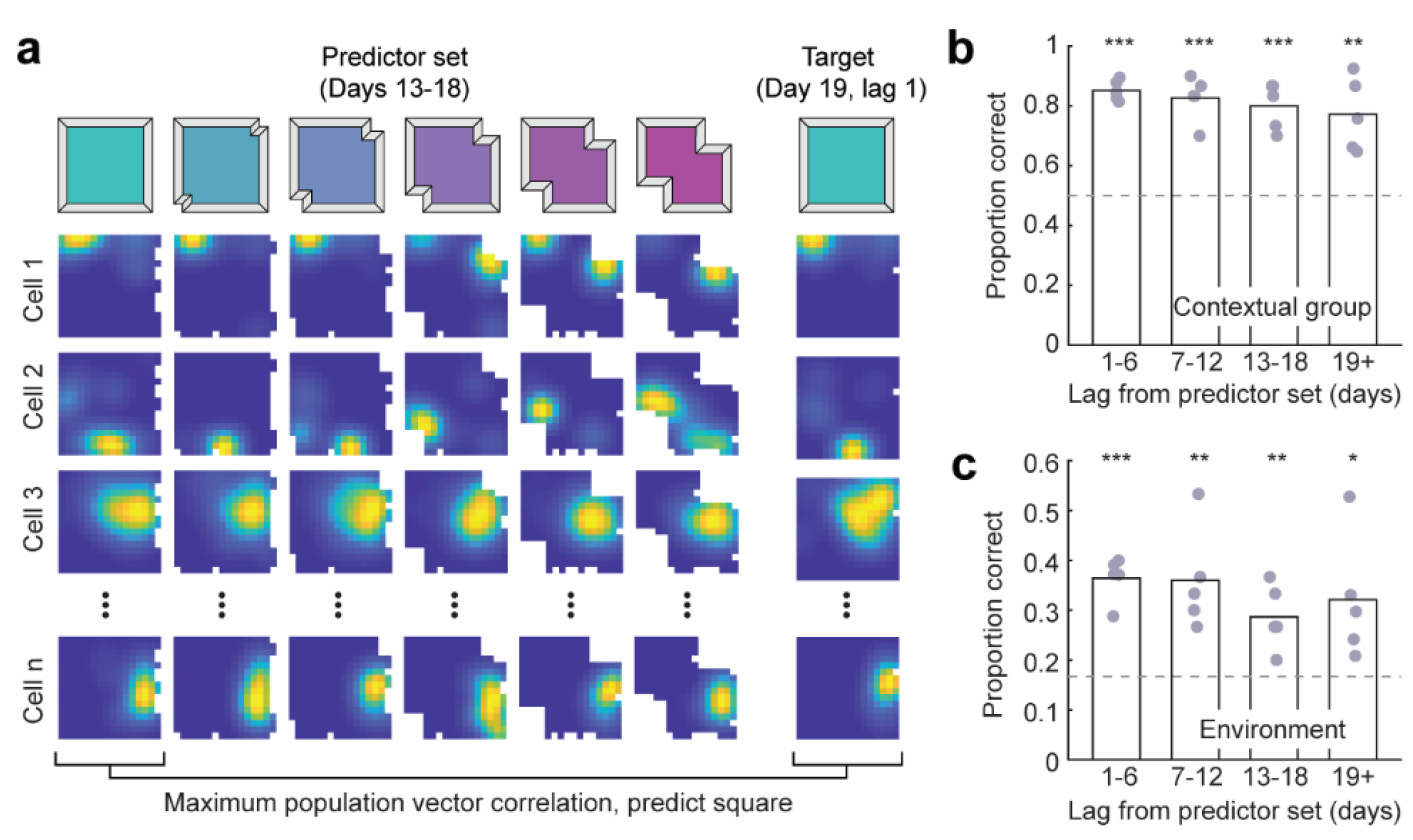
Drift preserves an accurate readout of contextual information across weeks. (**a**) To predict contextual group or environment identity, each target session was compared via population vector correlation to each environment in a 6-day contiguous predictor set morph sequence. Only cells which were tracked across all days of the predictor set and target session were included. The contextual properties associated with the highest population vector correlation to the target were then predicted for the target session. (**b**) Proportion correct for contextual group prediction as a function of lag between the predictor set and the target session (1-tailed t-test versus 0.5; (1-6): t(4) = 23.14, p = 1.03e-05; (7-12): t(4) = 9.61, p = 3.28e-04; (13-18): t(4) = 8.58, p = 5.07e-04; (19+): t(4) = 4.94, p = 3.90e-03; ANOVA between epochs: F(3,16) = 0.85, p = 0.488). (**c**) Proportion correct for environment identity prediction as a function of lag between the predictor set and the target session (1-tailed t-test versus 1/6; (1-6): t(4) = 9.94, p = 2.88e-04; (7-12): t(4) = 4.16, p = 7.05e-03; (13-18): t(4) = 4.13, p = 7.25e-03; (19+): t(4) = 2.76, p = 2.53e-02; ANOVA between epochs: F(3,16) = 0.82, p = 0.503). ***p < 0.001, **p < 0.01, *p < 0.05

The above results indicate that the representational differences which distinguished context were distinct from the representational differences which accompanied the passage of time at the level of the population. Such structure might arise from varying degrees of selectivity at the level of individual cells. Thus, we next examined the relationship between contextual representation and long-term stability at the level of individual cells. To quantify this relationship, we first characterized the extent to which the pattern of activity for each cell resembled an interpretable contextual code. To do so in a way which tolerates the heterogeneous but stereotyped responses we observed in our previous single-cell analysis, we again computed the mean squared error of the sigmoidal fit to the contextual RSM for each 6-day morph sequence for each cell as our (inverse) measure of contextual coding, as this fit could flexibly accommodate a range of interpretable dynamics (Fig. 1d; Fig. S2). To assay long-term stability, we computed the mean rate map correlation for the same environments across each pair of 6-day morph sequences. Finally, to ensure that separate data were used to compute each of these measures, we took the following approach. First, we computed the contextual RSM fit error for a given 6-day morph sequence for each cell tracked across this sequence (Fig. 5a). Next, we computed the stability between all remaining pairwise comparisons of morph sequences for each group, excluding the contextual fit sequence. We repeated this process for each contextual fit sequence, and combined the results across each lag between compared sequences. These two measures were negatively correlated at all lags (Fig. 5b), indicating that cells with a more interpretable contextual RSM (i.e. lower sigmoidal fit error) on a given sequence tended to have more stable rate maps across withheld sequences, even when withheld sequences were separated by weeks.

**Figure 5.**
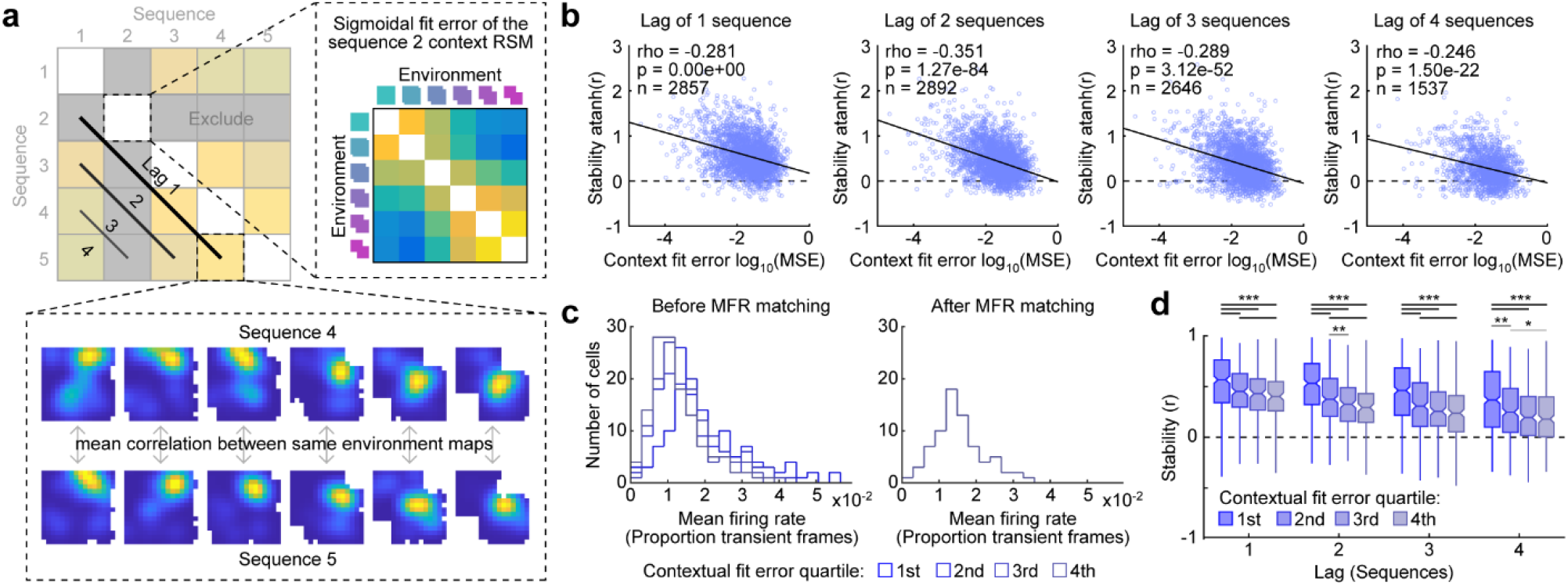
Interpretable contextual coding is associated with long-term stability at the level of individual cells. (**a**) Schematic for quantifying the relationship between contextual coding and long-term stability at the single-cell level. (**b**) Relationship between contextual RSM sigmoidal fit error and long-term stability at the single cell level, aggregated by the lag between the pairs of sequences from which long-term stability was measured. Dashed line indicates the least-squares fit. Outcome of the Spearman’s rank correlation noted. (**c**) Example of mean firing rate histograms for cells in each contextual fit error quartile before and after histogram matching by subsampling cells in each group. (**d**) Long-term stability at each lag for cells in each contextual fit error quartile, after subsampling cells in each group to match mean firing rate histograms. At all lags, cells in the 1^st^ quartile (i.e. cells with the lowest contextual fit error) had significantly greater long-term stability than cells in other quartiles, even after controlling for quartile differences in mean firing rates (Kruskal-Wallis test between quartiles: Lag of 1 sequence: Χ^2^(3) = 88.29, p = 5.10e-19; Lag of 2 sequences: Χ^2^(3) = 185.47, p = 5.80e-40; Lag of 3 sequences: Χ^2^(3) = 113.61, p = 1.83e-24; Lag of 4 sequences: Χ^2^(3) = 41.28, p = 5.71e-9; All significant Bonferroni-corrected post-hoc contrasts noted). ***p < 0.001, **p < 0.01, *p < 0.05

Notably, however, both of these measures were correlated with firing rate: cells which had higher firing rates tended to have lower average contextual fit errors (Spearman’s rank correlation: ρ = -0.345, p = ∼0.0, n = 3087) and higher rate map stability across sequences (Spearman’s rank correlation: Lag of 1 sequence: ρ = 0.275, p = ∼0.0, n = 3501; Lag of 2 sequences: ρ = 0.256, p = ∼0.0, n = 2761; Lag of 3 sequences: ρ = 0.201, p = 6.62e-23, n = 2379; Lag of 4 sequences: ρ = 0.165, p = 1.88e-12, n = 1797). To mitigate the influence of this potential confound when assessing the relationship of contextual coding and long-term stability, we took the following approach. First, we computed the contextual RSM fit error for a given 6-day morph sequence for each cell tracked across this sequence (Fig. 5a). Next, we divided these cells into four groups according to the quartile of their contextual fit error. We then randomly subsampled the cells in each of these four quartile groups to match the mean firing rate distributions across all four groups (Fig. 5c). Finally, we computed the stability between all remaining pairwise comparisons of morph sequences for each group, excluding the contextual fit sequence. We repeated this process for each contextual fit sequence, and combined the results across each lag between pairwise comparisons of sequences. This analysis revealed that cells with a more interpretable contextual RSM on a given sequence tended to have more stable rate maps across withheld sequences, even when controlling for covarying firing rate differences (Fig. 5d).

## Discussion

Here we recorded from CA1 in freely behaving mice over 32 days of experience in an adapted geometric morph paradigm. On a shorter within-sequence timescale the hippocampal representation resembled prior acute reports^8,9,24^, exhibiting sigmoidal population-level contextual similarity dynamics across the shapespace which were driven by heterogeneous but stereotyped single-cell patterns of activity. Characterizing the representational structure across all 32 days revealed that changes indicative of context were orthogonal in network space to changes which accompanied the passage of time. This specific structure was well described by a global drift model whereby drift is accrued independently of contextual identity. This structure enabled consistent readout of contextual information across a timescale of weeks even on the basis of a temporally-contiguous predictor set, suggesting that downstream contextual readout tuned to the representation at a particular time can successfully generalize across long timescales. Lastly, individual cells exhibited a positive relationship between interpretable contextual coding and long-term stability even when controlling for covarying differences in firing rate, suggesting that propensity to drift is heterogenous at the level of individual cells and linked to functional content. Together, these results demonstrate that the relative structure of the hippocampal representation of context is preserved despite continuous representational change on the timescale of weeks.

Recent results have provoked claims of representational drift at unexpected rates not only in the hippocampus^13–16^, but also in cortical sensorimotor regions^19,25,26^. In each of these cases, a claim of drift is made on the basis of critical assumptions about the form and content of the representation, specifically that the content of the representation is known and that the way this representation is encoded is also known. For example, for claims of drift within the hippocampus it is implicit in the analysis that the content of the representation is spatial and that the form of this representation is a population-wide rate code. While each of these assumptions are well motivated by prior work, there are alternative views that might accommodate the observed long-timescale dynamics without a claim of representational drift. Mounting cross-species experimental^27–31^ and theoretical^32–37^ work suggests that the hippocampal representation is fundamentally superspatial and includes a temporally-correlated dimension, either implicitly^33,37^or explicitly^32,35,36^. Moreover, it is known that the activity of hippocampal neurons is remarkably structured and coordinated on short timescales^38–41^, suggesting that a rate code (spatially-conditioned or not) may provide only a partial assay of representational content in this region. Because of these alternatives, the degree to which long-timescale changes to the hippocampal code reflect representational content versus drift remains open to debate, perhaps more so than in sensory regions where the coded content might be more completely understood.

The results we present here speak to this debate. Characterization of the population-level representational structure via dimensionality reduction revealed that changes in network activity which were indicative of spatial context were orthogonal to changes accompanying the passage of time, thus preserving the relative structure of contextual representation. While the changes accompanying the passage of time could reflect global drift independent of spatial context, this component could alternatively reflect a faithful reproduction of representational content which also varied on this timescale, for example representations of continuous experiences, predictive relationships, or time itself. Consistent with this, we observed a relationship between interpretable contextual coding and long-term stability, even when controlling for known firing rate covariates^42^. This link suggests that long-term *in*stability in spatial tuning properties might be driven by the representation of content beyond the spatial context. CA1 is known for genetic^43^, anatomical, and functional heterogeneity^44–46^ in the responses of individual cells, which is linked to biases in upstream inputs^47,48^. In addition to differences in their coding properties and content, these inputs also differ radically in their long-term stability^15,49^, providing a potential source for a link between representational content and long-term dynamics within CA1. This possibility thus motivates future work dissecting circuit-specific contributions to the representation of specific content and accompanying long-term dynamics.

Currently, it is unknown whether and how long-timescale changes attributed to drift are coordinated across the brain. One possibility is that drift is coordinated in such a way that incongruences between regions are continuously corrected^19^. On the other hand, here we show that information about the spatial context can be predicted with high accuracy across weeks despite ongoing representational changes, even on the basis of temporally-contiguous training data. This suggests that a downstream readout of contextual information trained at a particular moment in time will continue to generalize well on long timescales, even in the absence of continued coordination between the hippocampus and this hypothetical reader.

Here we leveraged the partial correlation between hippocampal maps of similar spaces to characterize the relationship between representational content and long-timescale dynamics. While necessary for this characterization, we are therefore limited in our ability to speak to ongoing changes in other portions of the representational statespace. Thus, while our results suggest that a global model in which long-timescale changes accrued independently of spatial context is most appropriate, we cannot rule out globally-inconsistent dynamics in other portions of the representational space.

In sum, our results demonstrate that the relative structure of contextual representation in CA1 is preserved despite continuous representational changes on the timescale of weeks. These results speak to an ongoing debate about representational content and drift throughout the brain, with fundamental implications for the nature of hippocampal coding. Finally, these results motivate future experiments dissecting the relationship between functional properties and long-term dynamics within and beyond the hippocampal formation.

## Methods

### Subjects

Naive male mice (C57Bl/6, Charles River) were housed in pairs on a 12-hour light/dark cycle at 22°C and 40% humidity with food and water ad libitum. All experiments were carried out during the light portion of the light/dark cycle, and in accordance with McGill University and Douglas Hospital Research Centre Animal Use and Care Committee (protocol #20157725) and with Canadian Institutes of Health Research guidelines.

### Surgeries

During all surgeries, mice were anesthetized via inhalation of a combination of oxygen and 5% Isoflurane before being transferred to the stereotaxic frame (David Kopf Instruments), where anesthesia was maintained via inhalation of oxygen and 0.5-2.5% Isoflurane for the duration of the surgery. Body temperature was maintained with a heating pad and eyes were hydrated with gel (Optixcare). Carprofen (10 ml kg^-1^) and saline (0.5 ml) were administered subcutaneously at the beginning of each surgery. Preparation for recordings involved three surgeries per mouse.

First, at the age of six to ten weeks, each mouse was transfected with a 350 nl injection of the calcium reporter GCaMP6f via the viral construct AAV5.CaMKII.GCaMP6f.WPRE.SV40 (CAMCA130) or AAV9.syn.GCaMP6f.WPRE.SV40 (all other mice). The original titre of the AAV9.syn.GCaMP6f.WPRE.SV40 construct, sourced from University of Pennsylvania Vector Core, was 3.26e14 GC-ml and was diluted in sterile PBS (1:1 AKCA170; 1:5 AKCA127; 1:10 AKCA130; 1:15 AKCA135) before surgical microinjection. The original titre of the AAV5.CaMKII.GCaMP6f.WPRE.SV40 construct, sourced from Addgene, was 2.3e13 GC-ml and was diluted in sterile PBS (1:3) before surgical microinjection.

One to three weeks post-injection, either a 1.8mm (AKCA127, AKCA130, AKCA135) or 0.5mm (AKCA170, CAMCA130) diameter gradient refractive index (GRIN) lens (Go!Foton) was implanted above dorsal CA1 (Referenced to bregma: ML = 2.0 mm, AP = -2.1 mm; Referenced to brain surface: DV = -1.35 mm). Implantation of the 1.8mm diameter GRIN lens required aspiration of intervening cortical tissue, while no aspiration was required for implantation of the 0.5mm diameter GRIN lens. Results observed using 1.8- or 0.5-mm diameter GRIN lenses were similar. In addition to the GRIN lens, two stainless steel screws were threaded into the skull above the contralateral hippocampus and prefrontal cortex to stabilize the implant. Dental cement (C&B Metabond) was applied to secure the GRIN lens and anchor screws to the skull. A silicone adhesive (Kwik-Sil, World Precision Instruments) was applied to protect the top surface of the GRIN lens until the next surgery.

One to three weeks after lens implantation, an aluminum baseplate was affixed via dental cement (C&B Metabond) to the skull of the mouse, which would later secure the miniaturized fluorescent endoscope (miniscope) in place during recording. The miniscope/baseplate was mounted to a stereotaxic arm for lowering above the implanted GRIN lens until the field of view contained visible cell segments and dental cement was applied to affix the baseplate to the skull. A polyoxymethylene cap with a metal nut weighing ∼3 g was affixed to the baseplate when the mice were not being recorded, to protect the baseplate and lens, as well as to simulate the weight of the miniscope.

After surgery, animals were continuously monitored until they recovered. For the initial three days after surgery mice were provided with a soft diet supplemented with Carprofen for pain management (MediGel CPF, ∼5mg kg^-1^ each day). Familiarization to both environments (while recording in the room A-associated environment to monitor imaging quality and habituate the mouse to recording) began 3 to 7 days following the baseplate surgery.

### Data acquisition

In vivo calcium videos were recorded with a UCLA miniscope (v3; miniscope.org) containing a monochrome CMOS imaging sensor (MT9V032C12STM, ON Semiconductor) connected to a custom data acquisition (DAQ) box (miniscope.org) with a lightweight, flexible coaxial cable. The DAQ was connected to a PC with a USB 3.0 SuperSpeed cable and controlled with Miniscope custom acquisition software (miniscope.org). The outgoing excitation LED was set to between 2-8% (∼0.05-0.2 mW), depending on the mouse to maximize signal quality with the minimum possible excitation light to mitigate the risk of photobleaching. Gain was adjusted to match the dynamic range of the recorded video to the fluctuations of the calcium signal for each recording to avoid saturation. Behavioral video data were recorded by a webcam mounted above the environment. Behavioral video recording parameters were adjusted such that only the red LED on the CMOS of the miniscope was visible. The DAQ simultaneously acquired behavioral and cellular imaging streams at 30 Hz as uncompressed avi files and all recorded frames were timestamped for post-hoc alignment.

All recording environments were constructed of a grey Lego base and black Lego bricks (Lego, Inc) according to the dimensions specified in the main text and supplemental figures. All external walls had a height of 22 cm. During recording, the environment was dimly lit by a nearby computer screen, which could serve as directional cue. During familiarization and all unrecorded experience the environments were well-lit by overhead room lighting. All sessions were 20 min, and only one session was recorded per day to avoid photobleaching. Following the recorded session, mice were returned to their home cage for 5 min, after which the unrecorded top-up experience in the familiar environments began. Mice were directly transported from one familiar environment to the other between unrecorded sessions. Each familiar environment session took place in a different neighboring room (e.g. room A:square environment, and room B:other familiar environment), with assignment of each familiar environment to a given room kept constant within each mouse and randomized across mice. All recordings took place in one of these rooms (e.g. room A). The two familiar environments were always recorded in the same order (e.g. square day n, other familiar environment day n+1) within each mouse, with the order randomized across mice. The order of the four morph environments was randomized for each sequence. The mouse was always placed in the same corner at the start of the session and was allowed to explore the environment for 15 to 30 s prior to the start of data acquisition. Following each session the environment was cleaned with disinfectant (Prevail).

### Data preprocessing

Calcium imaging data were preprocessed prior to analyses via a pipeline of open source MATLAB (MathWorks; version R2015a) functions to correct for motion artifacts^20^, segment cells and extract transients^21,22^. To extract the rising phase of transients from each filtered calcium trace, we proceeded as follows. First, we computed the derivative of the calcium signal, smoothed with a gaussian kernel with a standard deviation of 5 frames. Next, because calcium transients around the baseline can only be positive, we estimated the variance in the derivative of the smoothed calcium signal on the basis of noise via a half-normal distribution such that:

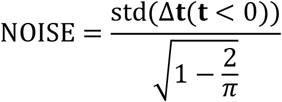

Where Δ**t** is the smoothed time-derivative of the median-subtracted calcium trace **t**. We then z-scored Δ**t** on the basis of this noise distribution. The final binarized rising-phase vector was then set to 1 whenever this z-scored Δ**t** vector exceeded 2.5, and 0 otherwise. This binary vector was treated as the *firing rate* in all further analyses. The motion-corrected calcium imaging data were manually inspected to ensure that motion correction was effective and did not introduce additional artifacts. Following this preprocessing pipeline, the spatial footprints of all cells were manually verified to remove lens artifacts. Position data were inferred from the onboard miniscope red LED offline following recording using a custom written MATLAB (MathWorks) script and were manually corrected if needed. Cells were tracked across sessions on the basis of prominent landmarks, their spatial footprints, and/or centroids^23^.

### Data analysis

All analyses were conducted using the binary vector of the rising phases of transients, treating this vector as if it were the firing rate of the cell (henceforth *firing rate*). Similar results were observed when the likelihood of spiking was inferred via a second-order autoregressive deconvolution^50^, instead of transient rising-phase extraction.

Rate maps were constructed by first binning the position data into pixels corresponding to a 2.5 cm x 2.5 cm grid of locations. Then the mean firing rate was computed for each pixel and then smoothed with a 4 cm standard deviation isometric Gaussian kernel. For all comparisons between rate maps, similarity was measured as the Pearson’s correlation between corresponding pixels. Individual cell contextual representational similarity matrices (RSMs) were thus computed as the Pearson’s correlation between pairwise rate map comparisons, resulting in a 6 × 6 matrix. To characterize individual cell contextual RSMs in various ways, each two-dimensional RSM was fit with a 5-parameter sigmoidal model as described by the two equations:

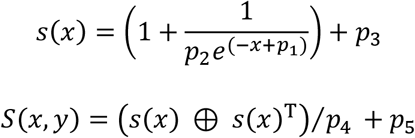

Where *s(x)* described a 1-dimensional sigmoidal function evaluated at *x, s(x)*^*T*^ denotes the transpose of this vector, ⊕ denotes the pairwise element multiplication of these two vectors [MATLAB’s *bsxfun*(@*times*,a,b)], and *p* is the vector of 5 free parameters to be fit by reducing the mean squared error between *S(x, y)* and the target contextual RSM. These parameters have intuitive interpretations: *p*_1_ determines the transition point for map similarity in the morph sequence, *p*_2_ determines the abruptness of the transition, *p*_3_ determines the asymmetry in similarity between halves of the shapespace, *p*_4_ determines the degree of sigmoidal modulation, and *p*_5_ determines the average overall similarity of the RSM. To evaluate the effectiveness of this description of contextual RSMs we also compared fit error for this model to a 5-parameter polynomial model (Fig. S2) which was described by:

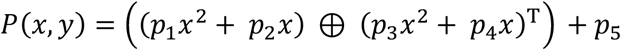

where again *p* is the vector of 5 free parameters to be fit by reducing the mean squared error between *P(x, y)* and the target contextual RSM. A two-dimensions embedding of contextual RSMs on the basis of their sigmoidal model fit parameters was created via Uniform Manifold Approximating and Projection (UMAP) with a Mahalanobis distance metric and a parameterization of k = 15 neighbors, minimum distance of 0.15, and 500 epochs. This embedding was clustered via k-medoids into 12 groups accepting the best of 100 replicates to avoid an initialization bias.

Within-sequence analyses summarized morph sequence dynamics with a transition plots (Fig. 2A), which captured the similarity of all 6 morph environment maps to the 2 familiar environment maps. To this end, rate map correlations between each environment and familiar rate maps at the beginning and end of the morph sequence were computed. Only comparisons between cells whose within-session split-half rate map correlation (SHC) exceed the 95^th^ percentile of a shuffled control for at least one of the compared sessions were included. The shuffled control was computed for each cell by randomly circularly shifting its firing rate vector relative to the position data by at least 30 sec and recomputing the SHC 1000 times to create the null distribution. To characterize these plots, the Fisher-transformed median values comparing the morph environments to each of the two familiar environments were both fit to minimize mean squared error with a 4-parameter sigmoid of the form

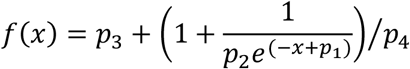

where *x* is the position of the current environment in the shapespace (arbitrarily chosen to range from 1 to 6), and parameters *p* determine the shape of the sigmoid. The intersection of these two sigmoids was taken as the measure of the transition point between familiar maps. The maximum absolute difference between these two sigmoids within the sampled shapespace was taken as the measure of maximum map decorrelation.

Two-dimensional embeddings of the population representation were computed and quantified as follows. First, we computed the mean rate map correlation across cells for each pairwise comparison of sections. Next, we transformed this correlation matrix into a distance matrix by computing one minus this correlation matrix, with the diagonal set to a distance of 0. Finally, this distance matrix was reduced to a two-dimensional embedding via Kruskal’s nonmetric multidimensional scaling, implemented by the MATLAB function mdscale with the default parameterizations including Kruskal’s normalized stress1 criterion. To estimate the context dimension within this embedding, the average difference separating neighboring-in-time familiar environment recordings was computed. To estimate the time dimension within this embedding, the average difference between the first and last recordings of each familiar environment was computed.

### Modeling CA1 dynamics

To simulate the representational structure of various drift dynamics, we created a rate map-based model. For ease of modeling, rate map binning was set in Lego coordinates, with the environment being 48 Lego pips wide (approximately equivalent to 38 cm).

First we created three populations of 48 × 48 pixel input rate maps which varied in their dynamics: a shape population (150 cells), a contextual group population (25 cells), and a drift population (300 cells). For shape inputs, a single pixels was selected as the field center for each cell. Next, for all sessions besides the square, if this pixel was in a row/column that was affected by the deformation then its location was rescaled to match the deformation. Half of geometry inputs were sensitive to the x-axis during deformations, with the other half sensitive to y-axis during deformations. For contextual group inputs, the three most square-like environments were assigned a single random pixel as the field center for those sessions. A different random pixel was then assigned for the remaining half of the shapespace.

Drift input rate maps were created differently depending on the modeled drift dynamics. In the case of global drift, a single pixel was randomly selected as the field center for each cell. Over consecutive days, this pixel was continuously shifted by a random amount on the range [-4, 4], inclusive, independently on both axes. For contextual group-specific drift dynamics, this drift accrued independently for environments in each half of the shapespace, such that over time each contextual group grew more dissimilar. For environment-specific drift dynamics, this drift accrued independently for every environment, leading all environments to become more dissimilar from one another over time. All input rate maps were then smoothed with an isotropic gaussian pixel with a standard deviation of 5 pixels, and normalized to have a maximum value of 1.

CA1 activity was then modeled as a rectified sum of random combinations of these inputs. For each CA1 cell, weights for a random 15 inputs were computed by assigning a random weight value on the range [0, 1] for each input and then normalizing all input weights to sum to 1. CA1 cell rate maps were then generated as the weighted sum of these inputs, thresholded at 75% of their maximum value and smoothed with an isotropic gaussian pixel with a standard deviation of 5 pixels. Resulting modeled CA1 cell rate maps tended to have 1-2 fields in each environment and appeared qualitatively similar to recorded rate maps.

### Histological validation of expression and recording targets

After experiments, animals were perfused to verify GRIN lens placement. Mice were deeply anesthetized and intracardially perfused with 4% paraformaldehyde in PBS. Brains were dissected and post-fixed with the same fixative. Coronal sections (50 μm) of the entire hippocampus were cut using a vibratome and sections were mounted directly on glass slides. Sections were split and half of all sections were stained for DAPI and mounted with Fluoromount-G (Southern Biotechnology) to localize GRIN lens placement and to evaluate virus expression. Due to the large imageable surface but restricted miniscope field of view (∼0.5 mm x ∼0.8 mm), we were unable to determine more specific localization of populations within the hippocampus for mice recorded with 1.8 mm lenses.

### Statistics and reproducibility

All statistical tests are noted where the corresponding results are reported throughout the main text and supplement. All tests were uncorrected 2-tailed tests unless otherwise noted. Z-values for nonparametric Wilcoxon tests were not estimated or reported for comparisons with fewer than 15 datapoints. Box plots portray the minimum and maximum (whiskers), upper and lower quartiles (boxes), and median (cinch).

### Code availability

All custom code written for reported analyses are publicly available at [insert Github link] or via request to the corresponding authors.

### Data availability

The complete dataset for all experiments are publicly available at [insert DRYAD link] or via request to the corresponding authors.

## Acknowledgements

We thank D. Aharoni for extensive guidance in using the UCLA miniscope. We thank J Quinn Lee for helpful feedback on prior versions of this manuscript. During this work ATK was supported by a McGill University Healthy Brains for Healthy Lives CFREF postdoctoral fellowship and a Natural Sciences and Engineering Research Council (NSERC) Banting postdoctoral fellowship. CAM was supported by a Fonds de Recherche du Québec – Santé (FRQS) postdoctoral fellowship. Funding was provided by the Canadian Institutes for Health Research (grants #367017 and #377074), the Natural Sciences and Engineering Research Council of Canada (Discovery grant #74105), the Canada Research Chairs Program, and the Brain Canada Foundation (Future Leaders in Canadian Brain Science).

## Author Contributions

ATK contributed to experimental design, surgeries, recordings, analysis of data, modeling, as well as drafting and revising the manuscript. CAM contributed to surgeries and histology. MPB contributed to experimental design, analysis of data, as well as drafting and revising the manuscript.

## Supplementary information for

**Supplementary Figure 1.**
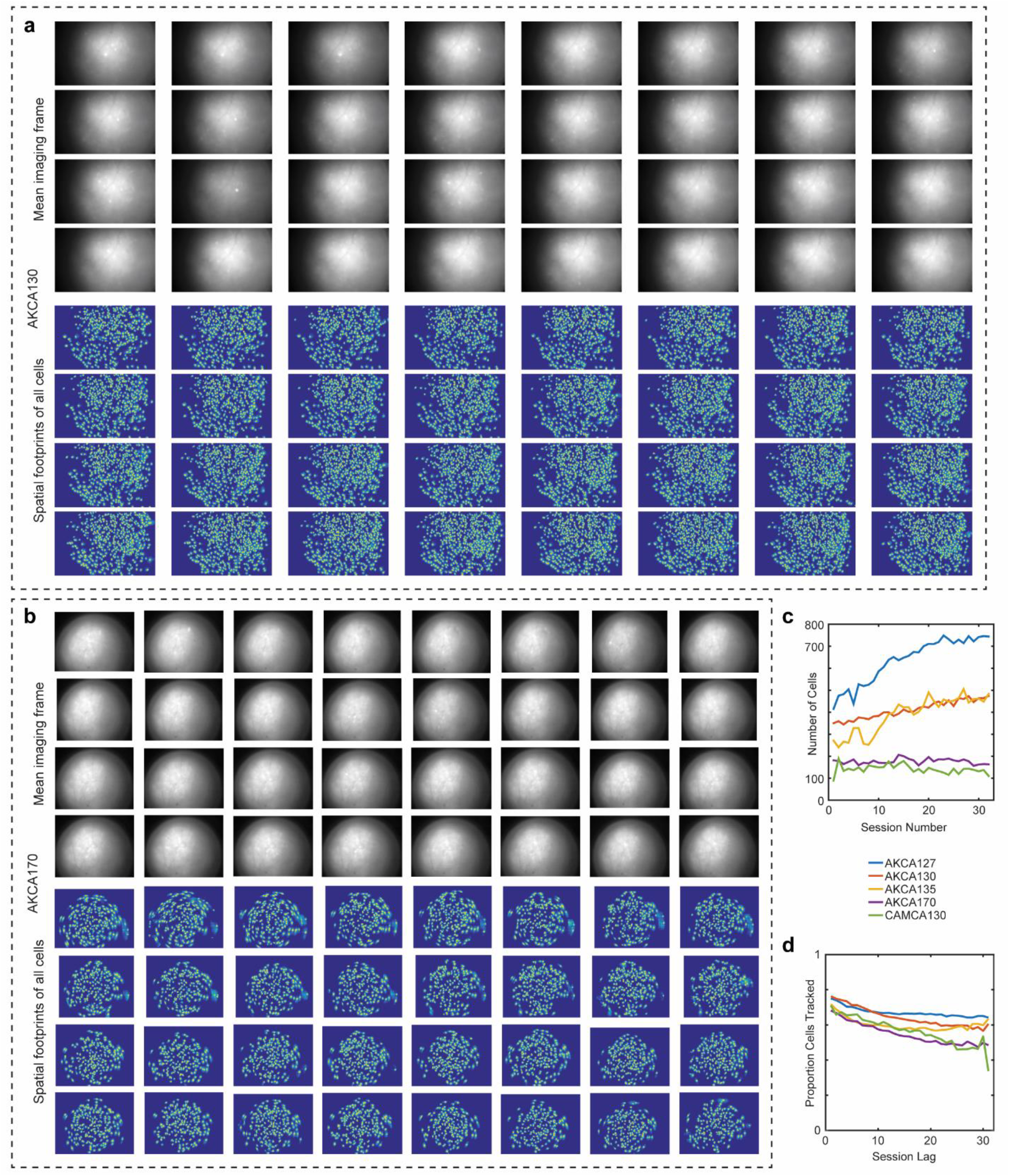
Imaging across 32 days of recording. Two example mice, AKCA130 (**a**) implanted with a 1.8 mm lens and AKCA170 (**b**) implanted with a 0.5 mm lens. Mean imaging frame normalized to maximum. Spatial footprints scaled and thresholded for ease of interpretation. Arranged chronologically. (**c**) Cell counts for each mouse across all 32 days. (**d**) Proportion of the population registered as a function of lag between sessions for each mouse. Normalized to the smaller population for each pairwise comparison.

**Supplementary Figure 2.**
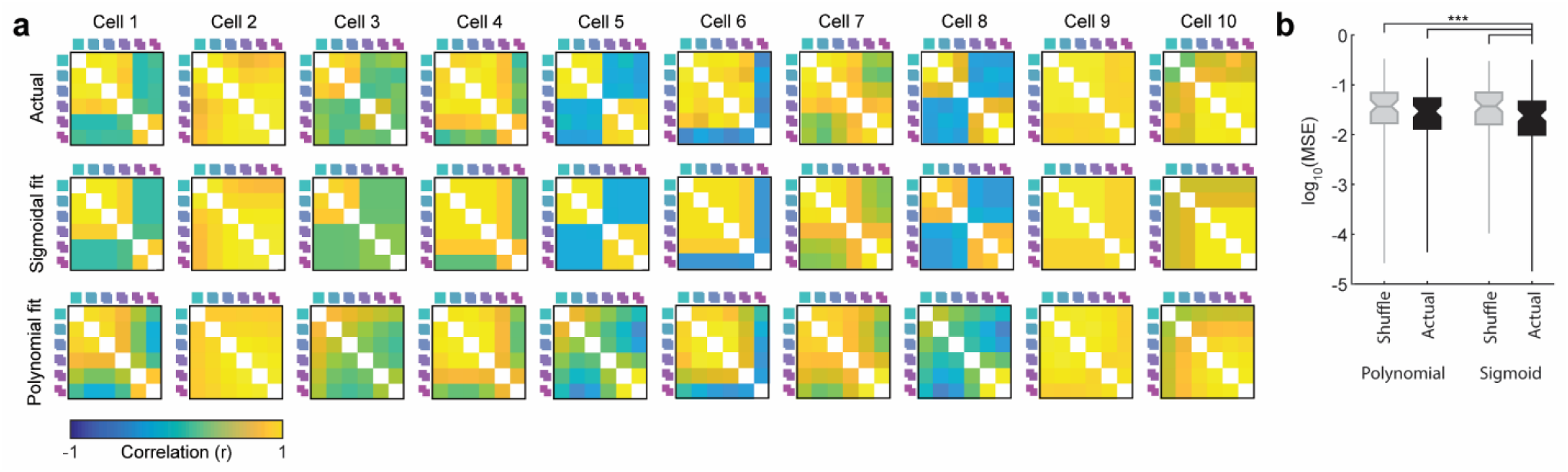
Contextual RSMs are well-described by a 5-parameter sigmoidal fit model. (**a**) Examples of actual contextual RSMs and the fits resulting from the 5-parameter sigmoidal model and the 5-parameter polynomial model for 10 cells (see *Methods*). (**b**) Mean squared error (MSE) for all models when fit to the actual RSMs and shuffled control RSMs in which environment identity was randomized for each cell (50 iterations). MSE was lowest for the actual RSMs fit by the sigmoidal model (Wilcoxon rank sum tests: Sigmoidal actual versus sigmoidal shuffle: W = 12203494721, Z = 21.28, p = 1.58e-100; Sigmoidal actual versus polynomial actual: W = 10020091, Z = 6.98, p = 2.90e-12; Sigmoidal actual versus polynomial shuffle: W = 12203887898, Z = 21.44, p = 5.44e-102).

**Supplementary Figure 3.**
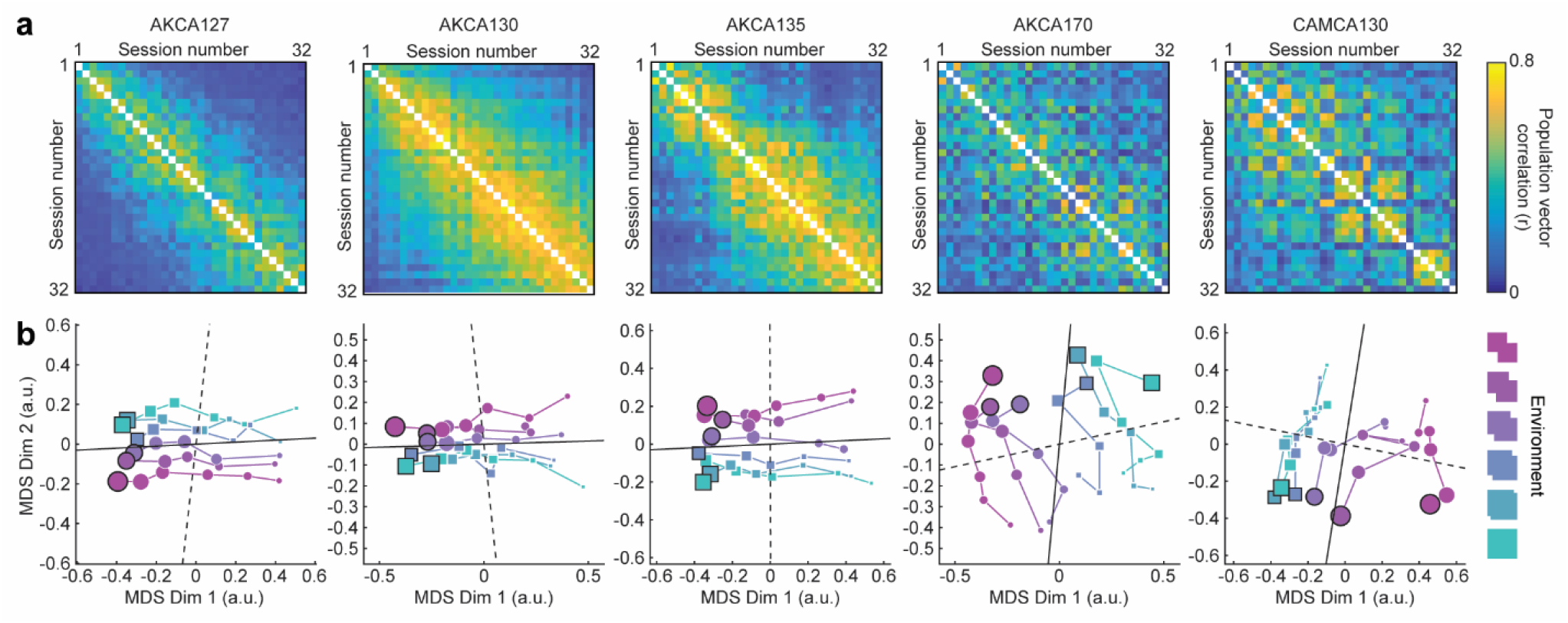
Assaying representational similarity via population vector correlations reveals a comparable structural embedding. (**a**) Population-level representational similarity matrices across all recording sessions for all five mice. Each RSM is computed by taking the population vector correlation between each pairwise comparison of sessions across all tracked cells. Mouse name at the top of each column. (**b**) Similarity structure when embedded in a two-dimensional space via nonmetric multidimensional scaling (MDS). Dot size indicates session number, with earliest sessions indicated by the smallest dots and latest sessions indicated by the largest dots. Final sessions outlined in black. Note that estimated drift (solid black line) and context (dashed black line) dimensions are nearly orthogonal in this embedding (see Methods).

